# Transcription-mediated organization of the replication initiation program across large genes sets up common fragile sites genome-wide

**DOI:** 10.1101/714717

**Authors:** Olivier Brison, Sami EL-Hilali, Dana Azar, Stéphane Koundrioukoff, Mélanie Schmidt, Viola Naehse-Kumpf, Yan Jaszczyszyn, Anne-Marie Lachages, Bernard Dutrillaux, Claude Thermes, Michelle Debatisse, Chun-Long Chen

## Abstract

Common Fragile Sites (CFSs) are chromosome regions prone to breakage under replication stress, known to drive chromosome rearrangements during oncogenesis. Most CFSs nest in large expressed genes, suggesting that transcription elicits their instability but the underlying mechanisms remained elusive. Analyses of genome-wide replication timing of human lymphoblasts here show that stress-induced delayed/under-replication is the hallmark of CFSs. Extensive genome-wide analyses of nascent transcripts, replication origin positioning and fork directionality reveal that 80% of CFSs nest in large transcribed domains poor in initiation events, thus replicated by long-traveling forks. In contrast to formation of sequence-dependent fork barriers or head-on transcription-replication conflicts, traveling-long in late S phase explains CFS replication features. We further show that transcription inhibition during the S phase, which excludes the setting of new replication origins, fails to rescue CFS stability. Altogether, results show that transcription-dependent suppression of initiation events delays replication of large gene body, committing them to instability.

## INTRODUCTION

The replication process should duplicate the entire genome once and only once at each cell cycle in normal conditions as well as under replication stress. In normal growth conditions, firing of tens of thousands of adequately distributed replication origins is needed to properly duplicate the human genome before mitotic onset. In cells submitted to mild replication stress, replication rate is supported by the recruitment of a large pool of extra-origins, an adaptation process called compensation ^1^. Conventional cytogenetic analyses have however shown that some regions of the genome, notably CFSs, display breaks in metaphase chromosomes of stressed cells, suggesting that replication completion recurrently fails in these regions ^2^. Consistently, CFSs are major drivers of genome instability and subsequent chromosome alterations associated with human diseases, notably oncogenesis ^3^. CFSs have long been associated with very large genes, in the order of the mega-base (Mb) ^4^, some of them behaving as tumour-suppressors ^5,6^. In addition, many very large genes have been associated with inherited diseases, such as neurodevelopmental and neuropsychiatric disorders ^7^.

Molecular mapping of CFSs in different human cell types has then extended the correlation between large genes and CFSs to genes over 300 kb long and, importantly, pointed out that CFSs display tissue specific instability ^8^. Strikingly, these genes and cognate CFSs are conserved in mouse ^4,8^ and chicken cells ^8,9^. Focussing on 5 large genes, a pioneer work has correlated the tissue-dependent expression of these genes to the instability of associated CFSs ^10^. ChIP-Seq of FANCD2, a factor binding preferentially CFSs from S-phase to mitosis upon replication stress ^11^, has recently confirmed genome-wide this co-localization of CFSs with large transcribed genes in human ^12,13^ and chicken ^9^ cells grown *in vitro*. Remarkably, the same correlation has been reached upon extensive mapping of copy number variations in large series of tumours ^14^. Therefore, it is now clear that transcription plays a major role in the setting of CFSs, which explains why different subsets of large genes are fragile in different cell types.

Two main models have been proposed to explain the susceptibility of large expressed genes to replication stress. The first one relies on the estimate that transcription of such genes takes more than one cell cycle, so that the machines of transcription and replication necessarily encounter during the S phase ^10^. When replication and transcription interfere, the most deleterious situation arises upon head-on encounters that favour formation and stabilization of R-loops, namely DNA/RNA hybrids resulting from annealing of the nascent transcript with the template DNA strand ^15,16^. It was therefore proposed that R-loops frequently form in the body of large genes, which delays fork progression and elicits CFS under-replication upon replication stress ^17^. A variant of this model proposes that stress-induced uncoupling of the replicative helicase from DNA polymerases gives rise to single-stranded DNA, which elicits formation of DNA secondary structures at particular sequences, notably at stretches of AT-dinucleotides. These structures would further delay polymerase progression, increasing the frequency of replication-transcription encounters ^18,19^.

The second model is based on molecular combing analyses of the distribution of initiation events along 3 CFSs, showing that the body of hosting genes is origin-poor in normal growth conditions and/or upon stress. Consequently, long-traveling forks emanating from flanking regions replicate those genes ^11^. The strong delay to replication completion that specifically occurs when such forks are slowed would elicit CFS instability. In this hypothesis, the role of transcription could be to chase origins from the gene body, as previously shown in various models ^20-24^. A direct support to this view was recently provided by replacement of the endogenous promoter of three large genes with promoters of various strengths, which has shown that the transcription level dictates the density of initiation events across the gene body ^25^. Many large transcribed genes however escape fragility indicating that, although important, transcription is not sufficient to set CFSs ^8^.

Replication timing is a second parameter often invoked to explain CFS susceptibility to replication stress. Indeed, late replication would increase the risk of leaving them under-replicated at the time of mitotic entry ^11^. But whether and how the replication timing impacts CFS stability remained poorly studied. Here we report Repli-Seq analysis of the replication program in human lymphoblasts grown in the absence or in the presence of aphidicolin (Aph), an inhibitor of replicative DNA polymerases used *in vitro* to destabilize CFSs ^2^. We identified regions displaying specific replication delay upon Aph treatment, resulting in under-replication. Over 80% of these delayed/under-replicated regions nest in chromosome domains transcribed continuously across at least 300 kb and poor in replication initiation events, thus replicated by long travelling forks. Strikingly, the orientation of those forks relative to transcription is neutral for establishment of delayed/under-replication. We further show that these regions correspond to a major class of CFSs. Noteworthy, inhibition of transcription in the S phase, a condition that suppress transcription-replication encounters but excludes origin resetting ^26^, does not rescue the instability of these CFSs. Altogether, our results demonstrate genome-wide that transcription-dependent segregation of initiation events out of the gene body generates long traveling forks in large transcribed domains, which elicits the replication timing delay responsible for CFS instability upon replication stress.

## RESULTS

### Fork slowing strongly affects the replication timing of specific loci

The impact of replication stress on the replication dynamics was quantified using the Repli-Seq technique (**Fig. 1a, Fig. S1a)**. We first determined the replication timing profile of human lymphoblastoid JEFF cells grown under normal conditions (Methods). The profiles were highly reproducible between 3 biological replicates (R>0.97, P<10^-15^) (**Fig. S1b**), and very similar to those previously reported for GM06990 and GM12878 cells, 2 other lymphoblastoid cell lines ^27^ (R>0.93, P<10^-15^) (**Fig. S1b**), which confirms that Repli-Seq is a robust technique and that the timing program is well conserved between different isolates of the same tissue ^28^. We then performed 2 biological replicates (R=0.95, P<10^-15^) (**Fig. S1b**) for JEFF cells treated prior to cell sorting with 600 nM of Aph for a total of 16 h (**Fig. 1a)**, conditions commonly used to induce breaks at CFSs. We computed the mean of reads per 50 kb window for the 3 replicates of control cells and the 2 replicates of Aph-treated cells, which allowed us to calculate a replication index (*RI*) defined as the Z score of the difference between the sum of reads per window in cells treated (Aph) or not (NT) with Aph (Δ_Aph-NT_, Methods). Negative (positive) *RI*s identified regions delayed/under-replicated (or not) upon stress.

**Figure 1.**
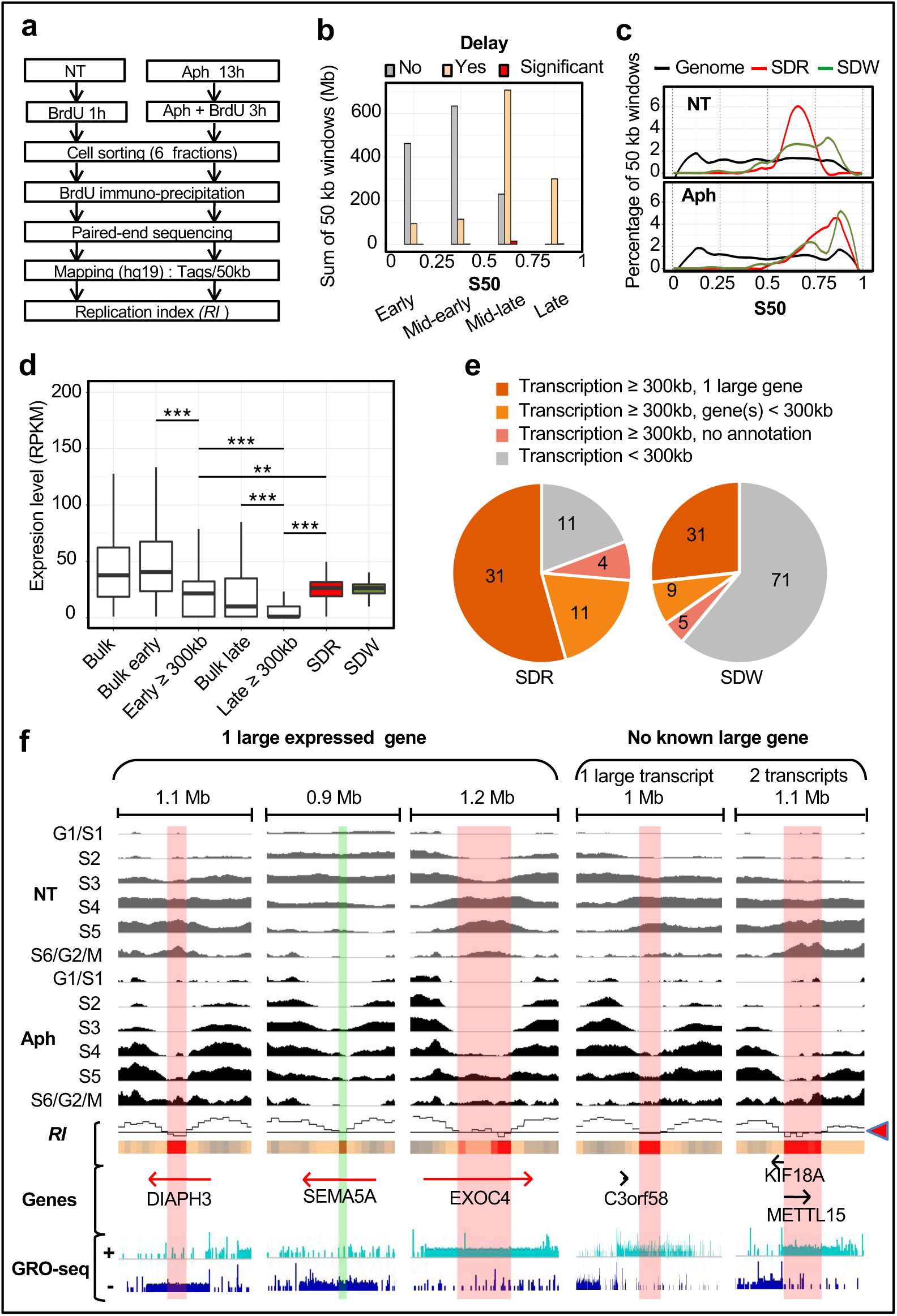
Genome-wide profiling of replication timing identifies SDRs/SDWs. **(a)** Scheme of Repli-Seq experiments performed with human lymphoblasts untreated (NT) and treated with 600 nm aphidicolin (Aph) for a total of 16 h. Cells were pulse labelled with BrdU as indicated then sorted in 6 fractions according to their DNA content. Purified BrdU labelled DNA was used to construct libraries for sequencing (Methods). The Replication Index (*RI*) was obtained at 50 kb resolution (Δ_Aph-NT_, Methods). **(b)** Histogram showing the sum of 50 kb windows (in Mb) displaying no delay (*RI*>0, grey, n=26558, 1328 Mb), low to moderate delay (−2<*RI*<0, orange, n=24343, 1217 Mb) or significant delay (*RI*<-2, red, *p*<0.05, n=330, 16.5 Mb) upon Aph treatment in the indicated classes of replication timing. The S50 is the moment in S phase, on a scale of 0 (Early) to 1 (Late), at which a given sequence has been replicated in 50% of the cells (Methods). **(c)** Percentage of 50 kb windows hosted in SDRs (red, n=57), isolated SDWs (green, n=116) and along the entire genome (black, n=56783) as a function of the S50 for non-treated cells (NT) and cells treated with aphidicolin (Aph). These percentages were computed within each bin (bin size=0.01) then Loess smoothed (alpha=0.25). For SDR, the replication timing values correspond to the mean of replication timings (S50) of all 50 kb windows enclosed in each SDR. **(d)** Boxplot showing the distribution of expression levels relative to gene replication timing. For genes over 50 kb long, the replication timing values correspond to the mean of replication timings (S50) of all 50 kb windows enclosed in each gene. The expression levels (RPKM, Reads Per Kilobase per Million of mapped reads, Method) were measured by GRO-Seq ^29^ for all expressed (Bulk, n=15204), for all genes replicated early (S50<0.5, n=13257) or late (S50≥0.5, n=1947), or the subset of genes≥300 kb free of SDR/SDW replicated early (n=315) or late (n=304) in the S phase, or for genes harbouring a SDR (n=54) or a SDW (n=39) (Kolmogorov–Smirnov test: ** *P*<0.01; *** *P*<0.001). **(e)** Pie-chart showing the numbers of SDRs or SDWs associated with the indicated classes of transcripts. **(f)** Representative examples of SDRs/SDWs identified upon analysis of the Repli-Seq profiles (50 kb sliding windows) normalized to the same background level as previously described (Methods). The enrichment in newly replicated DNA is shown for cells sorted at the indicated stages of the cell cycle following growth in the presence (Aph, black) or in the absence (NT, grey) of aphidicolin. The Replication Index profile (*RI*) is shown (the arrowhead points to the −2 threshold line) with the corresponding heat-map (colour coded as in **b**) below the *RI* profile. Examples of SDRs/SDWs nested within 1 long expressed gene (left panel, red arrows) are presented. SDRs/SDWs within regions where no large genes have been reported are shown (Right panel) together with potential candidate genes (black arrows). Examples of SDWs nested either in a large transcribed sequence (left) or a large expressed domain constituted of 2 adjacent transcribed sequences (right) are presented. Gene position and orientation are indicated together with GRO-Seq data showing read density on the Watson strand (+) and on the Crick strand (-). The extent of delayed/under-replicated sequences associated with SDRs (red vertical bars) and SDW (green vertical bar) is indicated. The genomic regions displayed are from left to right: chr13:61-61.1 Mb; chr5:8.8-9.7 Mb; chr7:132.8-134 Mb; chr3:143.4-144.4 Mb; chr11:27.7-28.9 Mb.

We then compared the *RI* to the replication timing using the S50, defined as the moment in S phase, on a scale from 0 to 1, at which a sequence has been replicated in 50% of the cells (Methods). We found that approximately half of the genome, essentially domains with a S50≥0.5 (mid-late and late), displays *RI*≤0 while the rest of the genome shows *RI*≥0. Importantly, we identified 330 highly delayed/under-replicated windows (*RI*≤-2, *p*<0.05), called below Significantly Delayed Windows (SDWs), out of which 314 nest in domains with a S50≥0.5 (**Fig. 1b)**. Genome-wide clustering of SDWs identified 57 regions hosting at least two SDWs separated by less than 250 kb (see Methods for the choice of this threshold), a distance significantly smaller than expected from random distribution (P < 10^-15^) (**Fig. S1c**, left panel). These regions, named Significantly Delayed Regions (SDRs), may display up to 16 SDWs and may extend over hundreds of kb (**Fig. S1c**, right panel; **Table S1)**. Together, SDRs enclose 214 SDWs, 116 remaining isolated (**Table S2**).

The timing profile of untreated cells shows that isolated SDWs distribute equally between mid-late (0.5≤S50≤0.75) and late (0.75≤S50≤1) replicating domains and are further delayed upon stress (**Fig. 1c, Table S2).** By contrast, 56 out of 57 SDRs (98%) nest in mid-late domains in untreated cells (**Fig. 1c, Table S1**). Upon stress, the SDRs however present much later S50 values and lower *RI* signals than all other mid-late domains and behave like late replicating domains (**Fig. 1c**). These results cannot be explained by a normalization bias in our data analysis since similar results were obtained with 3 different methods (**Fig. S1d**).

### SDRs/SDWs nest in large transcribed domains replicating in the second half of the S phase

We then looked for potential correlations between SDRs/SDWs and transcription features. Analysis of GRO-Seq data from untreated GM06990 lymphoblasts ^29^ showed that 31 SDRs nest within large (>300 kb) expressed genes (**Fig. 1d, 1e, 1f**). The human genome displays approximately 890 annotated genes larger than 300 kb, among which some 57% display a S50≥0.5, including those hosting SDRs. We observed that large genes, whatever their replication timing, are modestly transcribed compared to the bulk genes and that early replicating large genes are more transcribed than late replicating ones (**Fig. 1d**). Noticeably, the subset of genes hosting a SDR and/or a SDW displays an expression level resembling that of early replicating large genes, thus significantly higher than that of other late replicating large genes (*P*<10^-5^) (**Fig. 1d**).

We also found that 4 SDRs nest within regions displaying large GRO-Seq signal coming either from still non-annotated genes or from non-coding sequences, and 11 within long transcribed domains (>300 kb) harbouring 2-3 adjacent genes, which individual size may remain below 300 kb (**Fig. 1e, 1f, Table S1**). Together, 46 out of 57 SDRs (81%), named below transcription-associated SDRs (T-SDRs), nest in chromosome domains transcribed across at least 300 kb. Although the correlation appears less striking, 45 isolated SDWs (39%), named below T-SDWs, also nest in transcribed domains >300kb, which characteristics resemble those of the domains hosting SDRs (**Fig. 1e, 1f, Table S2**). Noticeably, 30 of the T-SDWs display at least one nearby window with a *RI* close to −2. These T-SDWs could therefore be false-negative T-SDRs resulting from our stringent *RI* cut-off (see *SEMA5* in **Fig. 1f**). T-SDRs/SDWs therefore mark moderately expressed large genes/domains, which body replicates in the second half of the S phase in normal conditions and displays strong retardation upon stress. Conversely, large transcribed genes, which replication completion occurs before S6/G2/M and non-transcribed large genes, even late replicating, do not show under-replication (**Fig. S1e**).

### The domains hosting T-SDRs/SDWs display low density in replication initiation events

We then analysed replication initiation in T-SDRs/SDWs and their flanking regions using data available for untreated GM06990 lymphoblasts. We first computed Bubble-Seq data ^30^ and found that over 80% of T-SDRs and T-SDWs, as well as their surrounding regions (several hundreds of kb to >1 Mb), are poor in initiation events when compared to genome-wide distribution (KS test *P*<10^-15^) (**Fig. 2a**). These conclusions were further supported and extended upon analysis of replication fork directionally (RFD) **(Fig. 2b**) determined by Okazaki fragment sequencing (OK-Seq) ^31^. In most cases, we observed that 2 major initiation zones flank the large transcribed genes hosting T-SDRs/SDWs. One generally overlaps the gene promoter while the second lies at variable distance from their 3’ end (**Fig. 2c, Fig. 3a**). Because of the size of the genes, unidirectional forks emanating from these zones travel across several hundreds of kb in order to complete replication of their body.

**Figure 2.**
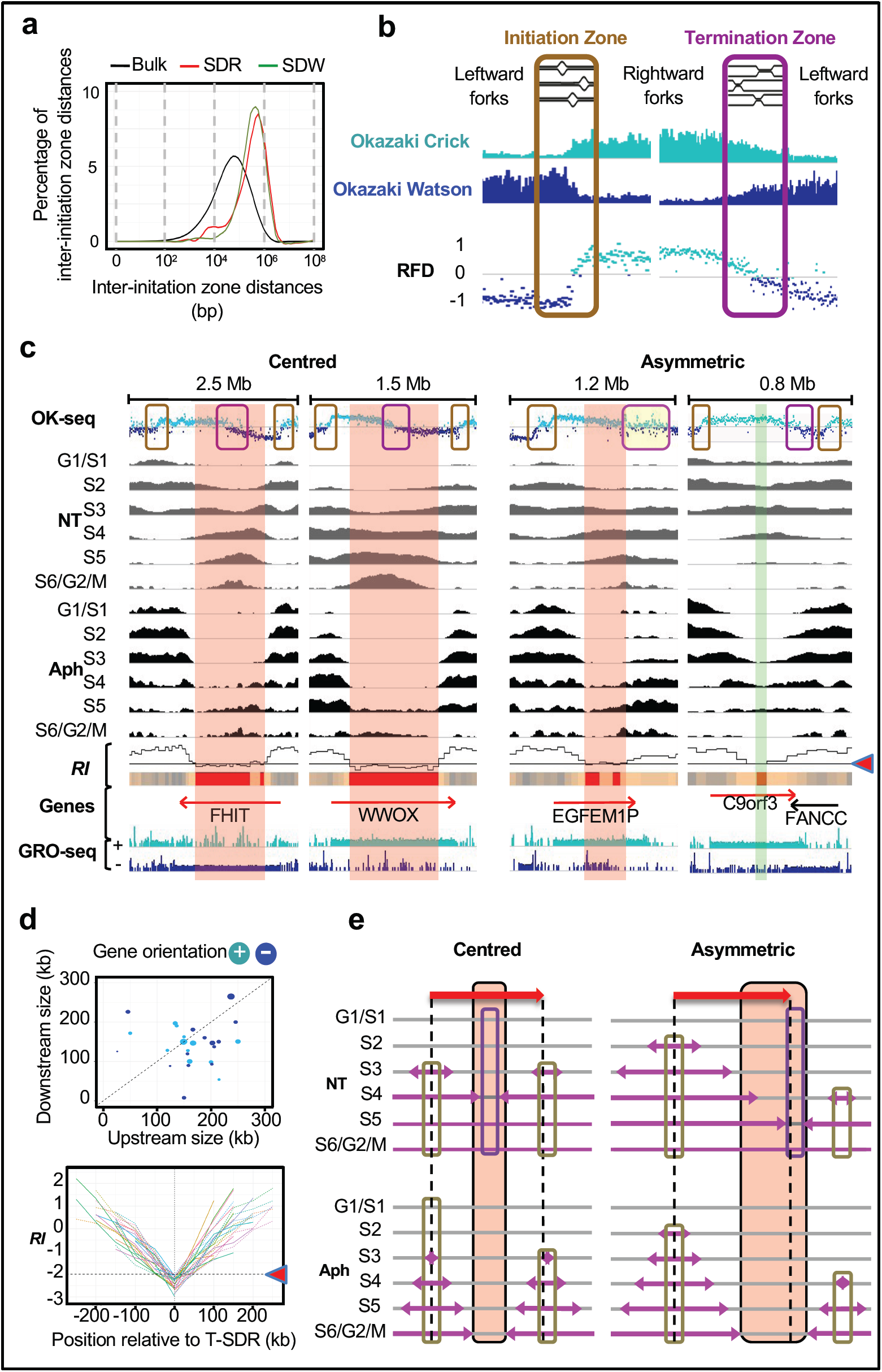
T-SDR/SDW localization relies on the properties of flanking replication initiation zones. **(a)** T-SDRs/SDWs are poor in initiation events. Percentage of inter-initiation zone distances identified by Bubble-Seq ^30^ for the inter-initiation zone regions overlapping SDRs (red), or SDW (green) compared to the distribution of inter-initiation zone distances along the genome (black). The percentages were computed as in Fig. 1c with bin size=0.1. **(b)** Principle of fork directionally (RFD) determination by Okazaki fragments sequencing (Ok-Seq). Okazaki fragments corresponding to rightward and leftward progression of replication forks map on the Crick (light blue) or Watson (dark blue) strands, respectively. Each point shows the RFD value, computed as the difference between the proportions of rightward and leftward moving forks in 1 kb windows. Upward (downward) transitions on the RFD profile correspond to replication initiation (termination) zone ^31^. **(c)** Representative examples of genes≥300 kb (red arrows) displaying a centred (left panels) or an asymmetric (right panels) termination zone on the OK-Seq profile of untreated cells. Repli-Seq, *RI* and GRO-Seq profiles, as well as T-SDRs/SDWs are as in Fig. 1f. The initiation and termination zones are highlighted by orange and purple boxes as in b, the yellow-filled box corresponds to a large zone where both initiation and termination events are taking place, leading to a complex RFD pattern. The genomic regions displayed are, from left to right chr3:59-61.5 Mb; chr16:77.9-79.5 Mb; chr3:167.6-168.9 Mb; chr9:97.4-98.2 Mb. **(d)** kinetics of *RI* decrease along sequences flanking the T-SDRs (n=46). Upper panel: Dot-plot showing the distances separating the transcription start site from the upstream end of the cognate T-SDR (Upstream size), and the transcription termination site from the downstream end of the T-SDR (Downstream size). Genes on the + or – strand are shown in light and dark blue, respectively and the sizes of the circle are proportional to the gene lengths. Note that the longest forks tend to come from the 5’ of the genes, independently of their orientation. Note that the longest forks tend to come from the 5’ of the genes, independently of their orientation. Lower panel: Each colour line shows the *RI* profile of the upstream and downstream (negative and positive values on the x-axis, respectively) regions flanking all individual T-SDR. The upstream and downstream extremities of each T-SDR have been both positioned at 0 on the x-axis and are shown in the same colour. The *RI* profiles show the progressive and regular decrease of *RI* values on both sides of the T-SDRs. Note that the *RI* decrease tends to be symmetrical at the global level as well as on both sides of individual T-SDRs. **(e)** Scheme depicting how initiation poor regions nested in large gene body elicits under-replication upon fork slowing. Pink arrows: replication forks progressing from the initiation zones in cells sorted in 6 fractions as above. Dotted lines show the limits of the genes. The genes, the initiation and termination zones and the T-SDRs/SDWs are represented as above. Left: two initiation zones with comparable firing times and efficiencies closely flank the 5’ and 3’ ends of a large active gene, the replication termination zone consequently maps at the centre of the gene, like the T-SDR/SDW. Note that the gene displays a symmetrical RFD profile on both sides of the T-SDR/SDW. Right: the large active gene is flanked by two initiation zones with different firing time and localization relative to the 5’ and 3’ ends of the gene, which position the termination zone out of the gene. Note the asymmetry of the T-SDR/SDW relative to the termination zone in untreated cells, with a shift toward the gene interior.

**Figure 3.**
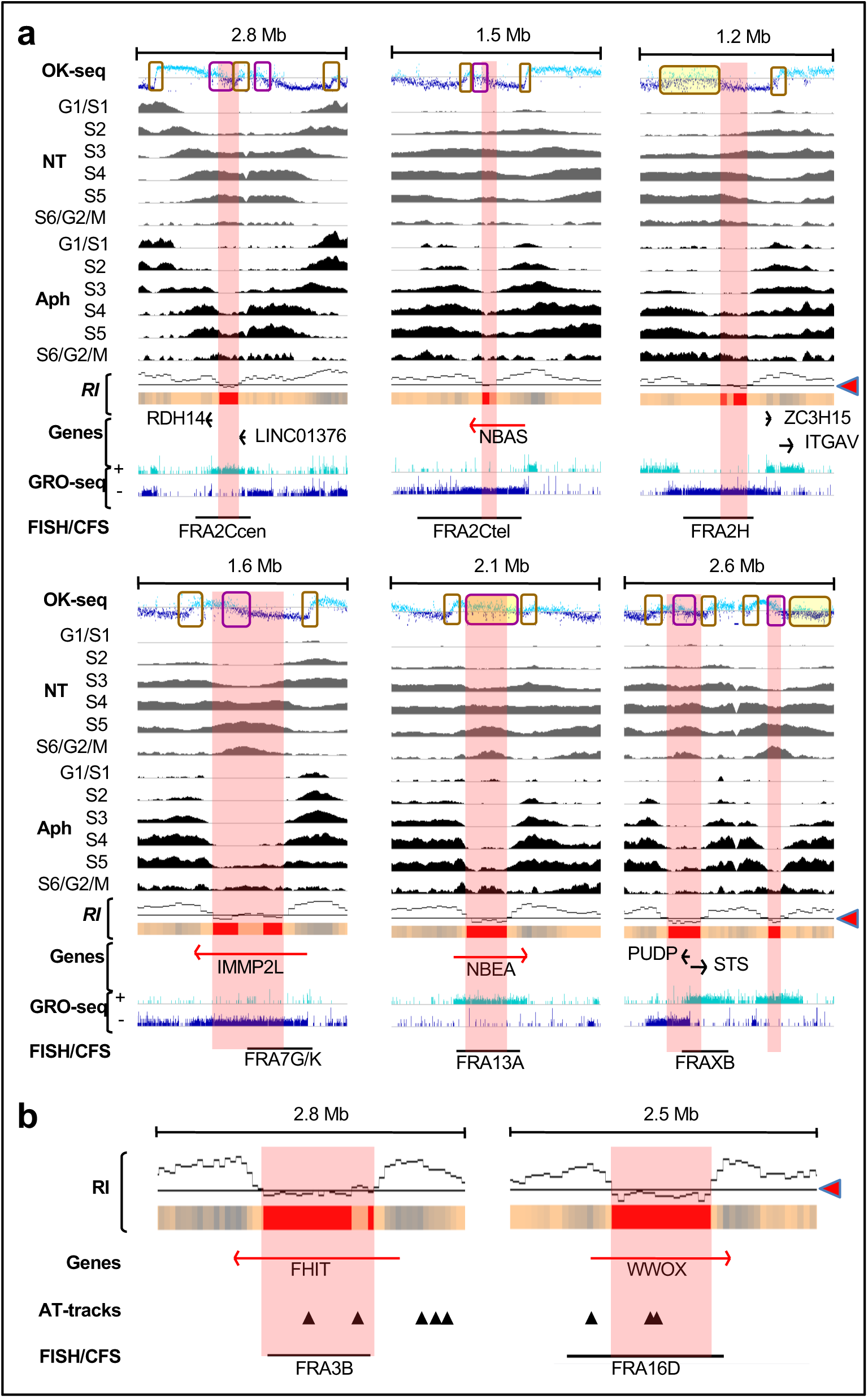
T-SDR/SDWs co-map with CFSs fine-mapped by FISH. **(a)** T-SDRs/SDWs positioning relative to the instable regions. OK-Seq, Repli-Seq and GRO-Seq profiles, and T-SDRs/SDWs (presented as in Fig. 2c) are shown along regions hosting the indicated genes/CFSs. These sites have been chosen because they display a break frequency ≥1% in normal lymphocytes ^32^, and have been fine-mapped by FISH (black bars) in lymphocytes ^19^. The T-SDRs/SDWs are included in, or at least partially overlap, the instable regions. The genomic regions displayed are: chr2:17.8-20.6 Mb; chr2:14.7-16.3 Mb; chr2:186.4-187.9 Mb; chr7:109.8-111.5 Mb; chr13:34.9-37 Mb; chrX:6.2-8.9 Mb. **(b)** Position of sequences susceptible to form secondary structures, notably AT-dinucleotide stretches, across the chromosome regions hosting FHIT/FRA3B and WWOX/FRA16D ^18^ (black arrowheads) relative to the *RI* profiles. The genes, *RI* profiles, T-SDRs/SDWs and instable regions mapped by FISH are presented as above. Note that the position of these sequences does not account for the regular and symmetrical *RI* decrease taking place over several 50 kb windows on both side of cognate T-SDRs.

We observed that the flanking initiation zones often fire early (**Fig. 2c, Fig. 3a**), sometimes very early in the S phase (**Fig. 2c**, *FHIT*), whether the cells were treated or not with Aph. However, the body of genes hosting T-SDRs/SDWs replicates in the second half of the S phase (**Fig. 1c**, NT), which shows that forks traveling long contribute to determine their replication timing. Note that, although poor in initiation events, the body of genes hosting T-SDRs/SDWs may display weak initiation zones, firing from S4 to S6, visible on the Repli-Seq profiles of cells treated with Aph. These initiation events tend to increase the *RI* locally, therefore help replication to proceed across large genes (**Fig 2c, Fig. 3a**). We conclude that initiation paucity is causal to under-replication of T-SDRs/SDWs.

### Fork to transcription direction is neutral relative to *RI* decrease

The OK-Seq profiles show that the termination zone associated with each couple of 5’ and 3’ initiation zones may lie at the centre of the large genes or in asymmetric position (**Fig. 2c, Fig. S2a, Fig. 3a**). Comparison of Repli-Seq and OK-Seq data reveals that the relative timing and efficiency of the flanking initiation zones, and their localisation relative to the transcribed domain impose the actual position of the termination zones. When the termination zones are centred in the transcribed domains, they are also centred in the cognate T-SDRs/SDWs. Noticeably, T-SDRs extend beyond the termination zone and largely overlap the terminal part of convergent forks, independently of their directionality relative to transcription (**Fig. 2c** left panel, **Fig. 3a**). When the termination zone is asymmetric, because the 5’ initiation zone fires generally first and more efficiently than the 3’ one, termination events take place closer to the 3’ of the genes than to its 5’, so that forks covering the longest distances progress co-directionally with transcription (**Fig. 2c** right panel, **Fig. 3a**). The inverse situation has been observed in only 2 cases (**Fig. S2a**). Since forks replicating the longest sequences are the most retarded upon Aph treatment, the T-SDRs extend in the domains replicated by the forks covering the longest distances in untreated cells, independently of their orientation relative to transcription (**Fig. 2c, S2a**).

In addition, we noticed that all T-SDRs/SDWs are flanked by regions along which the *RI* decreases progressively and regularly over 100-300 kb, independently of gene orientation and fork directionality (**Fig. 2c, 2d; S2 a, b**). In transcribed domains displaying centred termination zone, this decrease is nearly symmetrical on both sides of the corresponding T-SDRs/SDWs (**Fig. 2c, Fig. 3a**). When the termination zone is asymmetric, the slope of *RI* decrease may also be asymmetric, but remains progressive and regular on each side of the T-SDRs/SDWs (**Fig. 2c, S2a**). At the global level, the slopes of *RI* decrease are nearly similar for all T-SDRs/SDWs and remarkably independent of fork to transcription direction (lower panels of **Fig. 2d, S2b**). Our observations together show that the delay to replication completion imposed upon fork slowing mainly relies on the properties of the couple of initiation zones flanking each large transcribed domain. These conclusions are summarized in **Fig. 2e**.

### T-SDRs/SDWs co-map with CFSs

To determine whether T-SDRs/SDWs co-localize with CFSs, we mapped CFSs on R-banded chromosomes (Methods) from JEFF cells treated for 16 h with Aph 600 nM. Breaks were scored on 300 metaphase plates displaying a total of 320 breaks distributed among 59 loci (**Table S3**), of which 39 showed a break frequency ≥1% (**Fig. S3a**). Among these 59 loci, 58 co-map with CFSs previously localized in an extensive G-banding analysis of normal lymphocytes ^32^ (**Fig. S3a**), the last one (FRA3O) being described in fibroblasts and epithelial cells ^8^. Note that break frequencies at a given site may however vary between the different studies, which is not surprising since they also vary in normal lymphocytes from different individuals ^32^. Thank to this good concordance, we could use the large amount of data available from normal lymphocytes. Notably, the fine mapping of several sites by FISH ^19^ allowed us to precisely compare the position of instable regions to that of T-SDRs/SDWs. We found a very good concordance between the position of T-SDRs/SDWs and the most instable region of all 8 fined-mapped CFSs with a break frequency ≥1% in normal lymphocytes (**Fig. 3a, 3b**).

Forty-seven out of 59 chromosome bands displaying breaks in our conventional cytogenetic mapping (80%) contain T-SDRs and/or T-SDWs (**Table S3**), which generalizes the notion that T-SDRs/SDWs mark CFSs. Noticeably, the *FHIT* and *WWOX* genes that respectively host FRA3B and FRA16D, the 2 major sites in normal lymphocytes also highly instable in JEFF cells (**Table S3**), display the largest T-SDRs, approximately 800 kb each (**Fig. 2c; 3b**). Conversely, 38 T-SDRs (83%) and 32 T-SDWs (71%) nest in cytogenetic bands hosting CFSs mapped here and/or in normal lymphocytes ^32^ (**Table S1**). T-SDRs/SDWs are therefore the hallmark of a major category of CFSs.

Twelve CFSs mapped in JEFF cells by conventional cytogenetics remain free of T-SDRs/SDWs. Among them, FRA1E and FRA6C contain a large active gene (*DPYD* and *CDKAL1*, respectively) poor in initiation events and replicated in the second half of the S phase in untreated cells, suggesting that a few T-SDRs/SDWs may have escaped detection (**Fig. S3b**). For example, the minimum *RI* of *CDKAL1* is −1.83, slightly above the −2 cut-off, but raising the cut-off to 1.8 in order to include this gene resulted in a high number of false positive regions. Our method remains however highly reliable since we identified 39 out of 42 CFSs nested in large expressed domains (93%). Because of the large size of cytobands and in the absence of guides such as T-SDRs/SDWs or FISH mapping, the remaining sites cannot be further studied.

### Formation of sequence-dependent fork barriers does not account for *RI* decrease

Formation of cruciform and hairpin structures in regions enriched in stretches of AT-dinucleotide repeats has long been proposed to contribute to CFS instability because secondary structures could delay the progression of replication forks and eventually lead to under-replication ^18,19^. We therefore compared the *RI* profiles to the localisation of AT-dinucleotide repeats reported in the chromosome regions hosting FRA3B and FRA16D (**Fig. 3b**). Among them, 3/5 stretches of repeats found around *FHIT*/FRA3B and 1/3 found around *WWOX*/FRA16D, lie out or at the edge of the genes, in regions not or weakly delayed. Two stretches of repeats map in each 800 kb-long T-SDRs nested in *FHIT* and *WWOX* but their localization does not explain the extent of the SDRs. In addition, no repeat was found in the regions along which the *RI* decreases progressively and regularly down to −2 on each side of the *FHIT* and *WWOX* T-SDRs (**Fig. 3b**). We conclude that the distribution of the stretches of AT-dinucleotide repeats does not account for the *RI* profiles across these two genes.

### CFS instability does not rely on transcription-replication encounters

To directly evaluate the impact of transcription-replication encounters on CFS instability, we analysed JEFF cells soon enough after transcription inhibition to avoid resetting of origins along silenced genes. To this aim, we searched for conditions in which metaphase plates observed at the end of the experiment correspond to cells that were already in the S phase at the beginning of the experiment because building of new origins is prevented at this stage of the cell cycle ^26^. We turned transcription off with triptolide (Tpl), a drug that inhibits engagement of new RNA Pol II in expressed genes ^33^. In order to determine the shorter time of treatment able to clear genes of on-going transcription, we grew JEFF cells for different times in the presence of Tpl (**Fig. 4a**). We found that 3h of treatment are sufficient to clear genes of small or moderate size of nascent RNAs (**Fig. S4a**) but longer times are necessary to progressively clear the 1.5 Mb long *FHIT* gene (**Fig. 4a**). Noticeably, the clearing kinetics across *FHIT* (≈ 4 kb/min) agrees with genome-wide Pol II elongation rates previously measured in the body of genes poor in G/C and in exons ^34^, two features common to large genes hosting CFSs. We choose to pre-treat the cells for 5 h, the shorter time required to clear the *FHIT* body from most nascent RNAs. We also reduced the time of Aph treatment from 16 h to 7 h, which is sufficient to induce chromosome breaks at CFSs (**Fig. 4b** left panel), including FRA3B (**Fig. 4b** right panel). Note that compared to cells treated with Aph alone, the mitotic index was not strongly reduced in cells pre-treated for 5 h with Tpl alone then kept for 7 h in Tpl + Aph (Tpl+Aph) (**Fig. S4b**). We concluded that, although treatment with Tpl alone for 12 h impacts the mitotic index, the mitotic flow remains essentially dictated by Aph in cells grown in Tpl+Aph, which makes the two conditions easy to compare.

**Figure 4.**
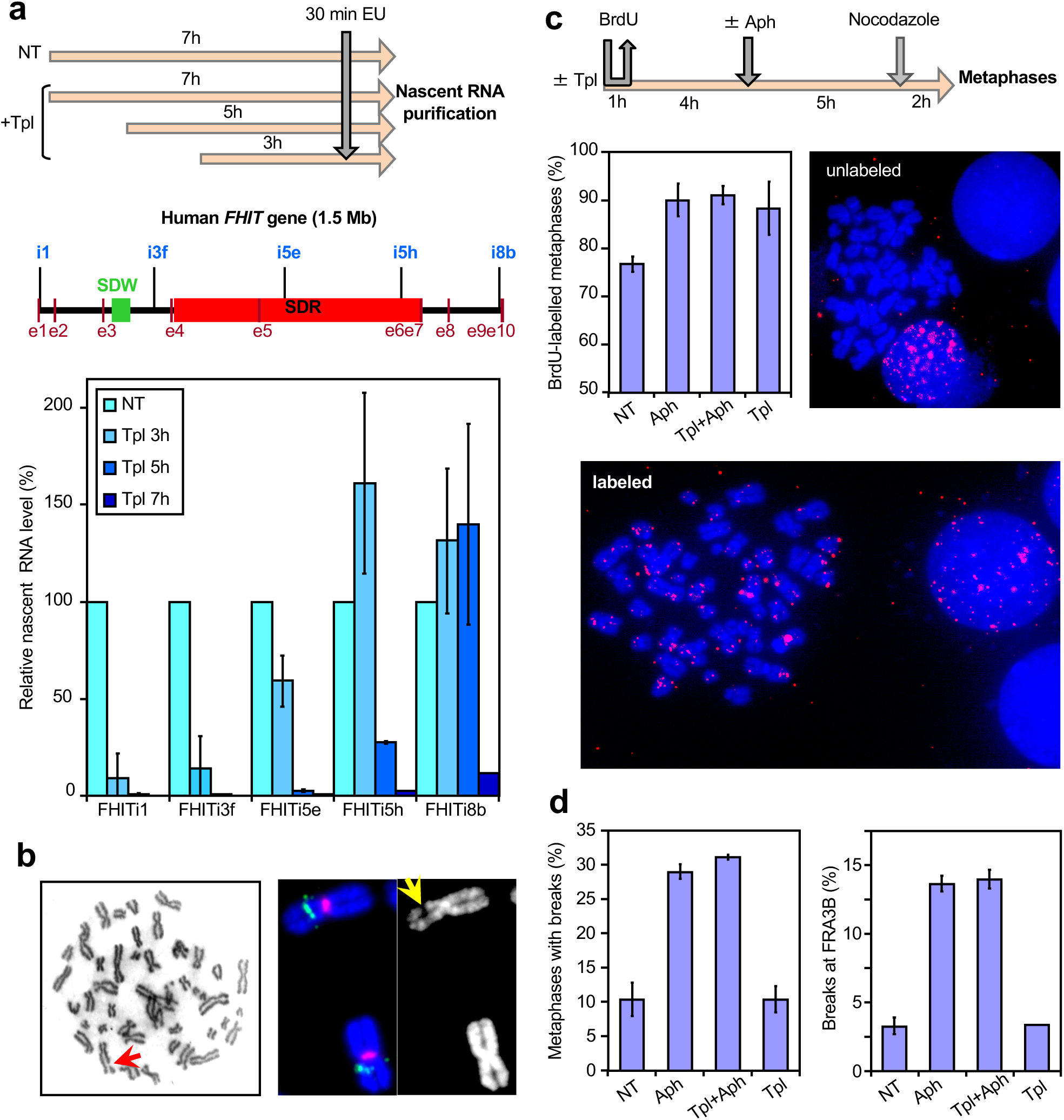
Short-term inhibition of transcription by triptolide does not impact CFS instability. **(a)** Determination of the minimal time permitting clearing of the *FHIT* gene from ongoing transcription. Upper panel: scheme of the experiments. Untreated cells (NT) and cells treated with triptolide 1 μM (Tpl) for the indicated periods of time were pulse labeled with EU for 30 min before recovery. Nascent RNAs were prepared by the Click-It method and quantified by RT-qPCR. Middle panel: map of the human *FHIT* gene with the position of exons (e1-e10), T-SDW/SDR, and intronic primer pairs (i1-i8b) used for quantification. Lower panel: quantification of nascent RNA along *FHIT* at the different times of Tpl treatment. **(b)** Examples of chromosome breaks in cells treated with Aph 600 nM for 7 h. Left panel: metaphase plate displaying a break (red arrow) on Giemsa stained chromosomes. Right panel: break at FRA3B (yellow arrow) visualized after DAPI staining and FISH with probes specific to the *FHIT* gene (green) and to the centromere of chromosome 3 (red). Contrast of DAPI stained chromosomes (rightmost panel) was enhanced to better show the break. **(c)** Upper panel: Scheme of the experiments: cells were treated with triptolide 1 μM and Aph 600 nM, alone or in combination, for the indicated times before metaphase preparation. In all conditions, the cells were pulse labeled for 1 h with BrdU 30 µM at the beginning of the experiments and treated for 2h with nocodazole 200 nM at the end of experiments to enrich the populations in metaphase cells. Lower panels: The fraction of BrdU labeled metaphases was scored after DAPI staining and immunofluorescence revelation with anti-BrdU antibodies. Examples of BrdU-labeled and unlabeled metaphases are shown. Note also the presence of labeled and unlabeled interphase nuclei. **(d)** Determination of the frequencies of total chromosome breaks (left panel) and breaks at FRA3B (right panel). Breaks were scored as in (**b**). Experiments shown in a, c and d were carried out twice and error bars represent standard deviation.

The percentage of BrdU-labelled metaphase plates recovered from cell populations pulse labelled at the beginning of the experiment was then determined (**Fig. 4c**). We found that ≈ 75 % of the metaphase plates were labelled when untreated cells were pulsed as indicated. In cells treated with Tpl alone, with Aph alone or with Tpl+Aph this percentage increases to ≈ 90 % (**Fig. 4c**). Counting total chromosome breaks in metaphase plates recovered from cell populations grown in each condition showed that Tpl alone, at least up to 12 h of treatment, does not increase the percentage of metaphase breaks relative to untreated cells while 7 h of Aph treatment increases this percentage by a factor of 3. Strikingly, Tpl fails to rescue chromosome breaks elicited by Aph treatment (**Fig. 4d, S4c**). These results were supported by FISH experiments with *FHIT*-specific probes showing that Tpl does not suppress Aph-induced breaks at *FHIT*/FRA3B (**Fig. 4d, S4c**). We conclude that transcription-replication encounters are not responsible for the instability of CFSs in these cells.

## DISCUSSION

Several mechanisms have been proposed to explain CFS instability, most of them postulating that CFS replication occurs in late S phase and is further delayed upon stress so that CFSs tend to remain under-replicated up to mitotic onset. Although consensual, this hypothesis stems from the study of only 3 sites ^11^ and a genome-wide analysis has failed to establish a link between CFS instability and replication features ^35^. The conclusions of the latter report are however questionable because the whole sequence of G-bands containing CFSs has been computed in this study, leading to a CFS map inconsistent with that established by cytogenetic analyses. Therefore, to what extent under-replication is common and specific to CFSs remained unclear. Here we searched blind for regions delayed/under-replicated in lymphoblasts treated with Aph, and studied the properties of identified loci. The main category of retarded loci corresponds to the so-called T-SDRs/SDWs that nest in large genes or adjacent genes behaving like a single large transcribed domain, which body replicates in the second half of the S phase and which transcription level is significantly higher than the bulk of genes with such replication timing. We then showed that almost all the T-SDRs and part of the T-SDWs overlap the most fragile region of fine mapped CFSs, or nest in cytogenetic bands hosting CFSs for those not mapped by FISH. Noticeably, our method proved to be highly sensitive and efficient for genome-wide mapping of CFSs since 32 out of 39 (82%) CFSs displaying a break frequency over 1% in conventional cytogenetic experiments were identified here at the sequence level, including 24 not yet fine-mapped.

CFS instability has been proposed to result from fork barriers raised by sequences prone to form secondary structures ^18,19,36^. At least 4 types of observations argue against this model; i) Analysis of the nucleotide sequence of several FISH-mapped CFSs has provided contrasted results regarding the presence of such roadblocks in a substantial proportion of the sites ^37^. In addition, recent mapping of CFSs genome-wide has failed to identify sequence features specific and common to CFSs ^9^. ii) The fact that CFS setting is tissue-dependent ^11^ strongly questions this hypothesis, at least in this simple form (see below). iii) We did not find abrupt drops of *RI* in the vicinity of particular sequences. Global analyses rather revealed that the *RI* decreases progressively and regularly across hundreds of kb and shows a near symmetrical slope on each side of the T-SDRs/SDWs. To account for these results, sequences forming fork barriers should distribute in a very particular manner, similar in all large fragile genes, which has never been reported. For example, although stretches of AT-dinucleotide repeats, a type of sequence proposed to raise fork barriers, lie in the T-SDRs of the *FHIT* and *WWOX* genes ^18,19^, their distribution does not account for the symmetrical 150-200 kb of progressive *RI* decrease flanking the 800 kb-long T-SDRs nested in either gene. In addition, molecular combing analyses of fork progressing across the *FHIT* gene in lymphoblasts treated or not with Aph did not show marks of fork barrier(s) ^38^. iv) Our data clearly correlate the slopes of *RI* decrease that culminates at T-SDRs/SDWs with the relative efficiency, firing time and localization of the initiation zones flanking the corresponding genes. Sequence features therefore have no major impact on CFS stability, except in particular genetic contexts in which resolution of secondary structures is impaired ^39-42^.

Other models rely on the association of CFSs with large expressed genes. One proposes that CFS instability is due to head-on encounter of the transcription and replication machines ^17,24^. However, as mentioned above, the *RI* decreases symmetrically across hundreds of kb on each side of the T-SDRs/SDWs, showing that the delay to replication completion is independent of fork direction relative to transcription. The finding that R-loops have short half-life ^43^ also argues against a major role of these structures in CFS instability, at least in cells proficient for factors ensuring their resolution, such as FANCD2 ^44^, which depletion enhances CFS instability ^45,46^. In agreement with this view, R-loop accumulation and replication fork stalling have been observed in the core of FRA16D in FANCD2 deficient cells but not in FANCD2 proficient cells ^47^, showing that cells cope with dynamic R-loops that form in the gene body ^48^. R-loops therefore do not appear as a major trigger of CFS instability, with the exception of particular genetic contexts.

Finally, we directly checked the impact of transcription-replication encounters on CFS instability. To this aim, we treated the cells with Tpl to clear the genes from the transcription machine and nascent RNA ^34^. The time course we choose allowed us to analyse metaphase plates coming from cells in which large genes were cleared of nascent RNA while already engaged in the S phase, a period during which setting of new origins is prevented by redundant pathways ^26^. We found that the frequency of Aph-induced breaks at CFSs is not affected under these conditions, again showing that transcription-replication encounters, whether head-on or head-to-tail, are not a significant source of CFS instability. Together, our results also strongly argue against the model proposing that fork progression is impaired by a combination of R-loops and secondary DNA structures ^18^. Note that our work strictly focuses on T-SDRs/SDWs and cognate late-replicating CFSs, our conclusions therefore concern neither early replicating fragile sites ^49,50^ nor other putative hard-to-replicate sequences ^18^.

Alternatively, the results presented here point to replication initiation paucity as a property common to CFSs nested in large expressed genes. Consistently, MCM7-^51^ and ORC2-^52^ ChIP-Seq experiments have revealed an under-representation of these components of the pre-replication complex ^26^ across genes associated with CFSs. Transcription-mediated chase of initiation events has been reported in the bulk genes of various organisms ^20-24^, which therefore shows that transcription shapes the replication initiation profile along transcribed sequences independently of their size and replication timing. However, fork slowing more drastically perturbs replication completion of loci replicated by long-traveling forks, a feature determined both by the size of the transcribed domain and by the degree of initiation paucity in the gene body, the latter property being controlled by the level of transcription ^25^.

In contrast to other models, the specific delay to replication completion elicited upon slowing of long traveling forks together with the relative properties of the initiation zones flanking the large genes readily account for the slope of *RI* decrease on both sides of the T-SDRs/SDWs. We also show here that those initiation zones often fire in the first half of the S phase, sometimes very early, indicating that forks traveling long are strongly involved in late replication completion of large gene body in unchallenged cells and in their under-replication upon slowing. We therefore conclude that the transcription program elicits initiation paucity, notably across large genes, generating long-traveling forks and subsequent late replication completion, which finally dictates the tissue-dependent landscape of CFSs.

## ACKNOWLEDGEMENTS

The 3 teams contributing to the work (M.D., CL.C., C.T.) have been supported by the Fondation pour la Recherche Médicale (FRM) (program DBI20131228560). M.D.’s team is also supported by the Institut National du Cancer (INCa) (subvention AAP INCa 2017 - PLBIO17-194). CL.C.’ team is also supported by the ATIP-Avenir program and by the Plan Cancer program. S. E-H. was supported by the FRM program. D.A. was supported by a fellowship from the Ligue Nationale Contre le Cancer. The authors would like to acknowledge the Imaging and Cytometry Platform (UMS 3655 CNRS / US 23 INSERM) of Gustave Roussy Cancer Campus for assistance with cell sorting, and the High-throughput sequencing facility of I2BC for its sequencing and bioinformatics expertise.

## AUTHOR CONTRIBUTION

M.D., CL.C. and C.T. conceived the project and analysed the results. M.D. and CL.C. wrote the paper. O. B. and S. E-H. contributed to writing and preparation of figures and tables. C.T. provided critical revision. O.B. contributed to- and directed D.A., M.S. and AM. L. bench work, and analysed the results. S. E-H. and CL. C. designed and performed bioinformatics and statistical analyses. D.A., S.K., M.S., V. N-K. and AM. L. contributed to biological experiments, including cell culture, cell sorting, immunoprecipitation of BrdU-labelled DNA and molecular cytogenetics. Y.J. did the Repli-Seq libraries and the sequencing. B.D. did the conventional cytogenetics mapping of CFSs in JEFF cells.

## COMPETING FINANCIAL INTERESTS

The authors declare no competing financial interest

## METHODS

### Cell culture

JEFF cells (human B lymphocytes immortalized with Epstein-Barr virus) were grown as previously described ^38^. Aph and Tpl were obtained from Merck (A0781 and T3652, respectively).

### Repli-Seq experiment

The technique was essentially as described by Hansen and coworkers ^27^. Exponentially growing lymphoblastoid cells (∼200×10^6^), were pulse-labelled with 50 μM BrdU (Merck, B-5002) prior to recovery. Untreated cells were pulse labeled for 1 h and Aph treated cells (600 nM, for a total of 16 h) were labeled for the last 3 h (Fig. 1a). Cells were then fixed in 70% ethanol and incubated overnight at 4°C in the presence of 15 μg/ml Hoescht 33342 (ThermoFisher Scientific, H3570). Cells were re-suspended in 1X PBS and sorted in six fractions at a time by flow cytometry based on their DNA content using a BD Biosciences INFLUX cell sorter. The first fraction contains cells in the second half of the G1 phase plus very early S (called G1/S1); the rest of the S phase was divided into four fractions (S2 to S5); and the last fraction contains cells in very late S, G2 and M phases (S6/G2/M) (Fig. S1a). To check the fractionation quality, the post-sorted cells, already stained with Hoescht 33342, were directly re-analyzed by flow cytometry. Cells were incubated overnight in lysis buffer (50 mM Tris-HCl pH8, 10 mM EDTA, 0.1% SDS, 50 μg/mL RNAse A, 100 μg/mL Proteinase K). DNA was purified from the lysates by phenol extraction. Four µg of genomic DNA was fragmented to a mean size of 500 bp on a Covaris S220 intrument (peak power: 140, duty factor: 10%, cycle/burst: 200, time: 80 seconds). Fragmented DNA was treated with the End-Repair module (New England BioLabs, E6050) and the A-Tailing module (New England BioLabs, E6053), according to the manufacturer recommendations. Illumina Truseq indexed adapters were ligated on the resulting fragments, using the Quick Ligation module (New England BioLabs, E6056), according to the manufacturer recommendations. Following heat denaturation, BrdU-DNA was isolated by immunoprecipitation using an anti-BrdU monoclonal antibody (BD Biosciences, 347580). Immunoprecipitated fragments were amplified for 10 cycles using the KAPA Hifi DNA polymerase (KAPABiosystems, KK2502) and the resulting libraries were purified with AMPure XP beads (Beckman Coulter, A63881). Illumina libraries were pooled and sequenced on a NextSeq500 instrument on Paired-end 2×43 or 2×75 bases, using a NextSeq 500/550 High Output Kit v2.

### Repli-Seq data processing

The Repli-Seq data were demultiplexed using the distribution of CASAVA software (CASAVA-1.8.2 bcl2fastq2 v2.18.12). Illumina adapters were removed using Cutadapt-1.15, keeping only reads with a minimal length of 10 nucleotides. The reads were mapped on the human genome (Hg19), the chicken genome (galGal4, BrdU label DNA as positive control) and the salmon genome (GCF_000233375.1_ICSASG_v2, unlabelled DNA as negative control) using bwa-0.6.2-r126. The mapped data were then processed as previous described ^53^, with the following modifications. The PCR duplicates were removed by Picard tools (http://broadinstitute.github.io/picard) and the paired-end reads mapped properly to a unique position of the genome were kept for downstream analysis. The sequence reads located within the regions likely resulting from the sequencing hotspots (defined as the 0,5% windows with the highest amount of reads within 200bp windows) were also removed. The read density (*D*_*w,Si*_) was them computed for each 50 kb non-overlapping windows (w) for each sample *S*_*i*_ corresponding to different *S* phase fractions (i = 1–6) as well as for the control sample of cells within entire *S* phase named *S0* (density *D*_*w,S0*_). The background levels were then estimated as previously described ^53^. The replication timing, S50, defined as the moment in S phase, on a scale of 0 (Early) to 1 (Late), at which a given sequence has been replicated in 50% of the cells, was computed by linear interpolation of enrichment values in the 6 compartments of the *S* phase. When a region was not significantly enriched in all 6 *S*_*i*_ periods, no S50 value was attributed (∼5% of genome regions, most located close to telomeres or centromeres). The S50 values of biological replicates were strongly correlated to each other (R > 0.95, P < 10^−15^) (Fig. 1c; Fig. S1). The mean S50 values of the biological replicates were therefore used in downstream analyses. The raw sequencing data together with the processing data are available in GEO with accession number GSE134709. The raw Repli-Seq data of other lymphoblastoïd cells (GM06990 & GM12878) were downloaded from the Encode project (http://genome.ucsc.edu/cgi-bin/hgFileUi?db=hg19&g=wgEncodeUwRepliSeq) and the S50 were computed.

### SDR Identification

To identify the genomic loci, the replication of which is specifically delayed upon aphidicolin treatment, we computed the difference between the amount of newly replicated DNA measured by Repli-Seq in cells treated with aphidicolin (Aph) and that obtained in non-treated cells (NT). A replication Index (*RI*) was defined as the Z-score computed on the Δ(Aph – NT) by using all 50 kb window along the human genome, where 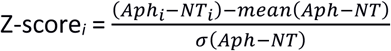, measuring the difference between the Aph and NT signals summed over the six periods ∑_*j*=1,6_ (*Aph*_*j*_ – *NT*_*j*_), where the mean and *σ* were computed by all ∑_*j*=1,6_(*Aph*_*j*_ – *NT*_*j*_) along the genome. The windows with a *DI* < −2 (n=330, 0.54% of genome) were defined as windows with significantly delay (*p*<0.05), called significant delay windows (SDWs). To limit the false positive results, the windows with too low or too high amount of reads were removed and only the windows with average coverage, mean (NT, Aph), between 20 and 40 after normalization, were retained. The SDWs were frequently close each other forming clusters (Fig. S1). Hence, the close windows (n≥2) passing the filtering process and separated by a distance<250 kb (maximum distance between adjacent SDWs located within fine-mapped CFSs) were then merged as a significantly delayed region (SDR).

### Metadata analyses

The RNA-Seq data of GM12878 cells from ENCOE project (GEO: GSM758559, GSM758559_hg19_wgEncodeCshlLongRnaSeqGm12878CellPapGeneGencV7.gtf) were used. Annotation of genes was retrieved according to gencode V7. The level of transcription was calculated in RPKM (Reads Per Kilobase per Million mapped reads) for each protein coding gene. The genes with RPKM > 0.0001 for both biological replicates were kept and the mean values of the two replicates were used. Raw GRO-Seq data of GM12878 cells generated in ^29^ (GEO: GSM1480326) were used for measuring the transcriptional level for each gene. GRO-seq read densities on the corresponding strand were calculated along all 1 kb non-overlapping windows along each gene (gencode v7) and the median value was then computed for each gene. Inter-origin distances in GM06990 lymphoblastoïd cells were calculated by using replication origins identified by Bubble-Seq ^30^. Only the Bubble-Seq origins identified in at least 2 biological replicates were retained in the analysis. The replication fork directionality (RFD) data of GM06990 cells determined by sequencing of Okazaki fragments (OK-Seq) ^31^ as well as the replication initiation zones, termination zones and regions replicated by unidirectional replication forks were used (SRA: SRP065949). The Repli-Seq, Ok-Seq, Bubble-Seq and GRO-Seq data were analyzed using custom scripts written in Python (v2.7.9) and R (v3.4.4) and the data were visualized with the Integrative Genomics Viewer (IGV)^54^.

### Cytogenetic analysis

The populations were enriched in metaphasic cells by 2 h of treatment with 200 nM nocodazole (Merck, M1404) prior to cell recovery. Total breaks were counted on metaphase plates stained with Giemsa (Prolabo) without pre-treatments to obtain a homogeneous staining of the chromosomes. Preparations were then de-stained and treated to reveal R-bands as previously described ^55 8^. Preparations were re-stained with Giemsa, and the breaks previously detected were localized relative to the bands. FISH on metaphases, Giemsa counter-staining, and immunofluorescence revelation of BrdU labeled DNA or FHIT-specific FISH probes were carried out as previously described ^38,55^. Chromosome 3 centromeric probe was from Aquarius Probes (LPE03R).

### Nascent RNAs isolation and quantification

Five-ethynyl-uridine (EU) (Life Technologies, E10345)) was added to cell culture medium (1 mM final) during the last 30 min of treatments (Fig. 4a). Total RNA was extracted with the miRNeasy kit (Qiagen, 217004) and nascent RNA was isolated using the Click-It Nascent RNA Capture kit (Invitrogen, 10365) and streptavidin coated magnetic beads (Dynabeads MyOne Streptavidin T1, Invitrogen 11754). After dissociation of bead-bound RNA by heating (70°C, 5 min) cDNA synthesis was carried out using the Superscript VILO cDNA Synthesis kit (Invitrogen, 11754). RNA/cDNA hybrids were then incubated for 5 min at 85°C and quantification was carried out by qPCR with specific primer pairs (Fig. 4 and S4; sequences of primers are available upon request).

## DATA availability

All sequencing files and processed count matrices were deposited with the Gene Expression Omnibus (GEO) under accession number GSE134709. Previously published accessions have been included in the Methods section where appropriate.

## Code availability

The computer codes and further processing data are available upon request to Chun-Long Chen.

## FIGURE LEGENDS

**Figure S1. Repli-Seq analysis of lymphoblasts treated or not with Aph.**

**(a)** Scheme of Repli-Seq experiments. Upper panel: Cells at 6 successive periods of the S phase G1/S1, S2, S3, S4, S5 and S6/G2/M were obtained by FACS sorting from asynchronous cell populations previously pulse labelled with BrdU. Lower panel: Schematic representation of forks progressing from an early-firing initiation zone. Evolving position of BrdU labelled DNA (red) is shown across the replicon at different periods of the S phase. Purification and sequencing of BrdU labelled DNA at the 6 steps of the S phase allow genome-wide reconstitution of replication fork dynamics. **(b)** Correlation coefficients (Pearson’s *r*) of replication timing (S50) between untreated GM06990 and GM12878 lymphoblastoid cells, untreated JEFF cells (3 replicates, NT1, NT2 and NT3) and JEFF cells treated with Aph (2 replicates, Aph1 and Aph2). Colour intensity and the size of the circle are proportional to the correlation coefficients. **(c)** Clustering of SDW. Left panel: Boxplot comparing the distance separating SDWs (n=330) upon random simulation (Control, black) or upon analysis of our data (green). SDWs separated by less than 250 kb were merged as SDR (Methods). Right panel: Histogram showing the distribution of SDRs according to their SDW content. **(d)** Comparison of different normalisation methods. Upper panel: *RI* profiles (as in Fig. 1f) at 3 loci upon normalization to the same total number of reads (i.e 15M), by adjusting the background to the same level (background) or by the level of mitochondrial DNA (mtDNA). Lower panel: Veen diagram showing the number of overlapping SDRs with the different normalization methods. **(e)** Representative examples of large genes without T-SDR/SDW (displayed as in Fig. 2c): Late replicating and not (or very weakly) transcribed (Left), early replicating and transcribed (right). The genomic regions displayed are from left to right: chr1:238,711,214-240,908,847; chr12:26,397,342-27,126,35.

**Figure S2. Relationships between replication forks and transcription directionality**

**(a)** Asymmetric termination zones lying in 3’ of two large genes in untreated cells, so that forks progress head-on with transcription all along the gene in untreated cells and across most of the gene in Aph-treated cells. Ok-Seq, Repli-Seq, *RI* and GRO-Seq profiles are shown as in Fig. 2c. Displayed genomic regions are chr2:148.2-149.6 Mb and chr11:30.7-32.5 Mb, from left to right. **(b)** Analysis of *RI* profiles along sequences flanking the T-SDWs (n=45, upper and lower panels: as in fig. 2d).

**Figure S3. Correlations between CFSs mapped by conventional cytogenetics and SDRs/SDWs**

**(a)** The thirty-nine CFSs with a break frequencies ≥1% mapped in JEFF cells are presented according to Mraseck *et al.*^32^ and ordered by decreasing break frequencies. Coloured bands contain at least one SDR (red) and/or SDW (green). CFSs identified by a white arrowhead are associated with T-SDRs or T-SDWs, FRA6C/H (green arrowhead) might be a false negative (see Fig. S3b). **(b)** Examples of expressed large genes instable upon Aph treatment but free of T-SDRs/SDWs that might be false negative. OK-Seq, Repli-Seq, *RI* and GRO-Seq profiles are presented as in Fig. 2c. Both genes are free of SDR/SDW, but the *RI* profiles show delayed regions close to −2, all along the very late domain hosting the gene (DPYD) or in the middle of the gene like in canonical CFSs (CDKAL1). Green vertical bars point to the latest windows, *ie* putative SDWs. The genomic regions displayed are chr1:96.4-99.4 Mb and chr6:20.1-21.8 Mb.

**Figure S4. Impact of Tpl on transcription, cell cycle progression and CFS instability**

(a) Maps of cyclophilin (*PPIB*) and *MYC* genes with positions of exons and primer pairs (upper panel) used to quantify nascent RNAs by RT-qPCR in the indicated conditions (lower panel) are shown as in Fig. 4a. **(b)** Microscopic counting of metaphase plates relative to total nuclei after Giemsa staining (left panel) allows determination of the mitotic index in each condition (right panel). **(c)** Representative examples of Giemsa stained metaphase plates from untreated cells, cells treated with Tpl alone or with Tpl+Aph (the red arrow points to a chromosome break). Note that chromosomes of Tpl-treated cells are over-condensed. Example of break at FRA3B (yellow arrow) visualized after DAPI staining and FISH with probes specific to *FHIT* (green) and centromere of chromosome 3 (red) after Tpl+Aph treatment. Contrast of DAPI stained chromosomes (right panels) was enhanced to clearly show the break (compare to the unbroken second allele).

