## Supplementary figures and images for "Transcription-mediated organization of the replication initiation program across large genes sets up common fragile sites genome-wide"

### Supplementary Figures

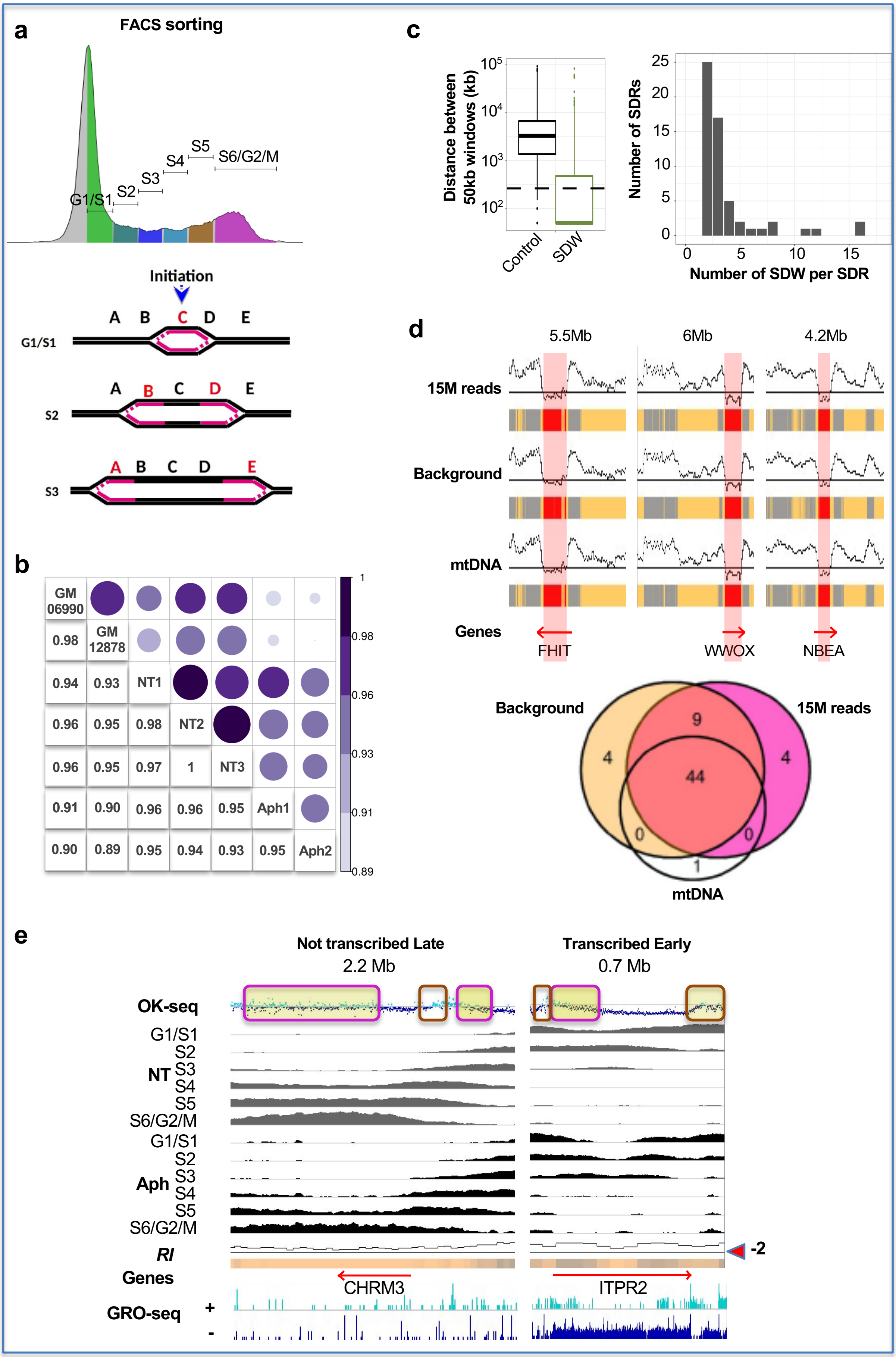

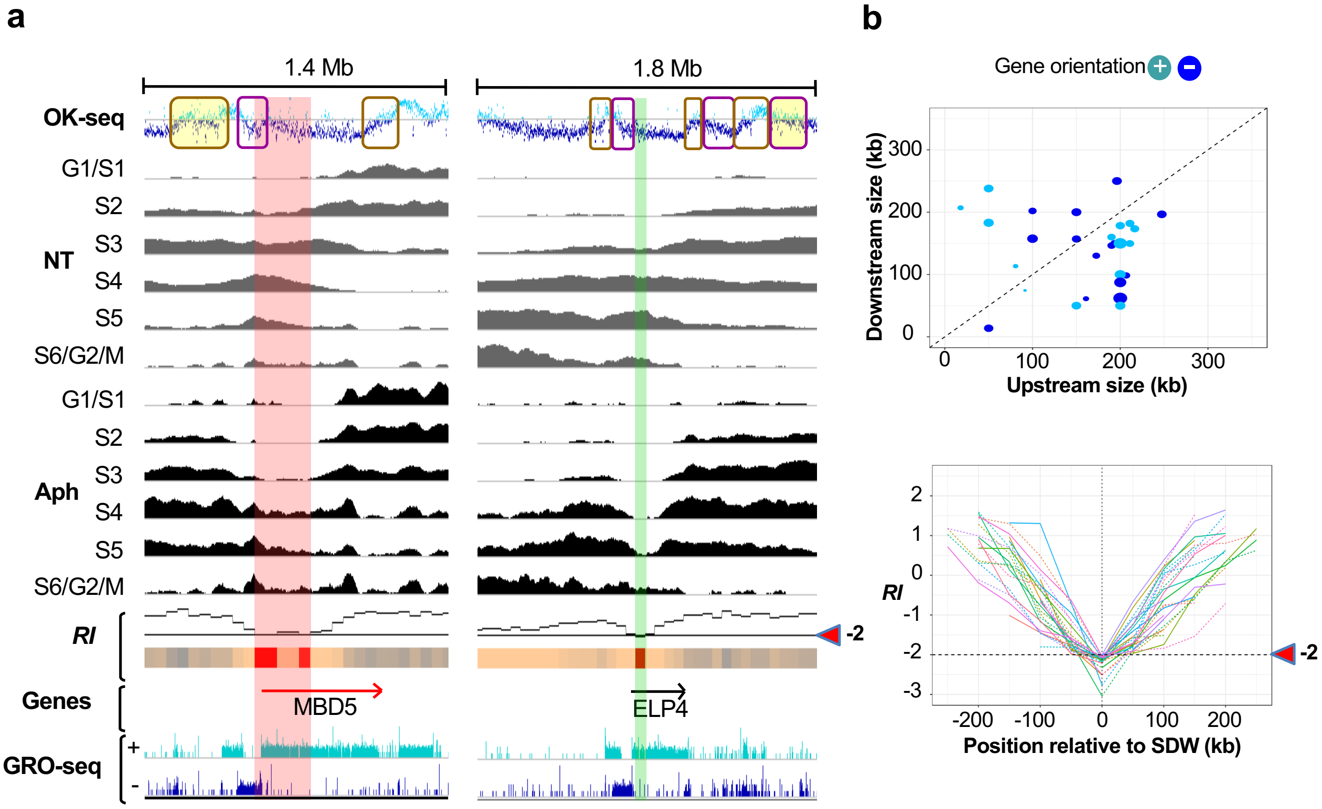

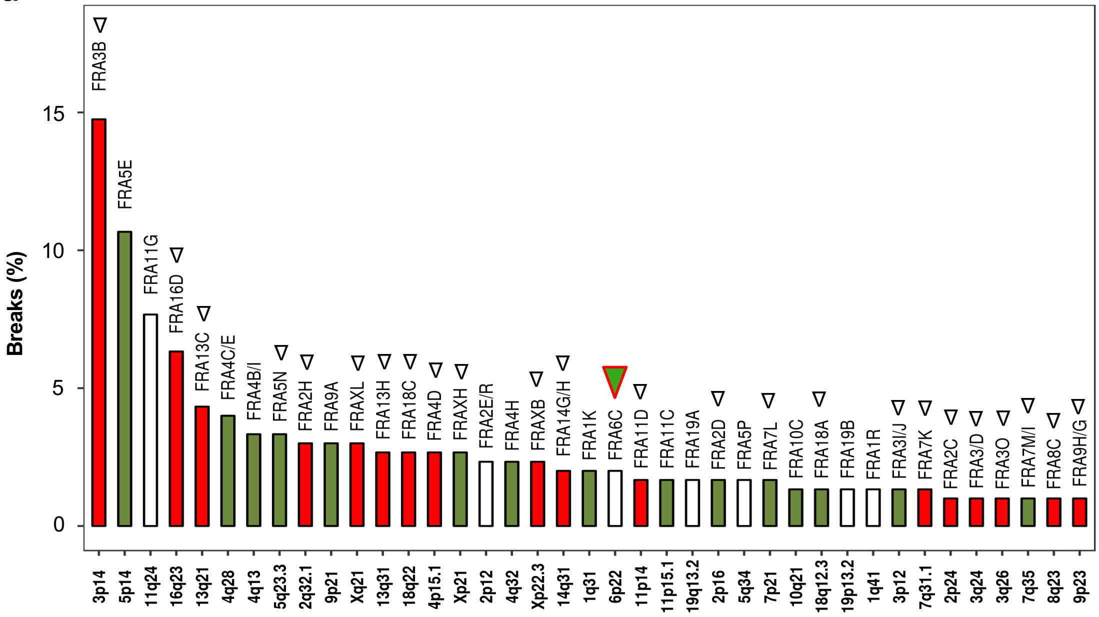

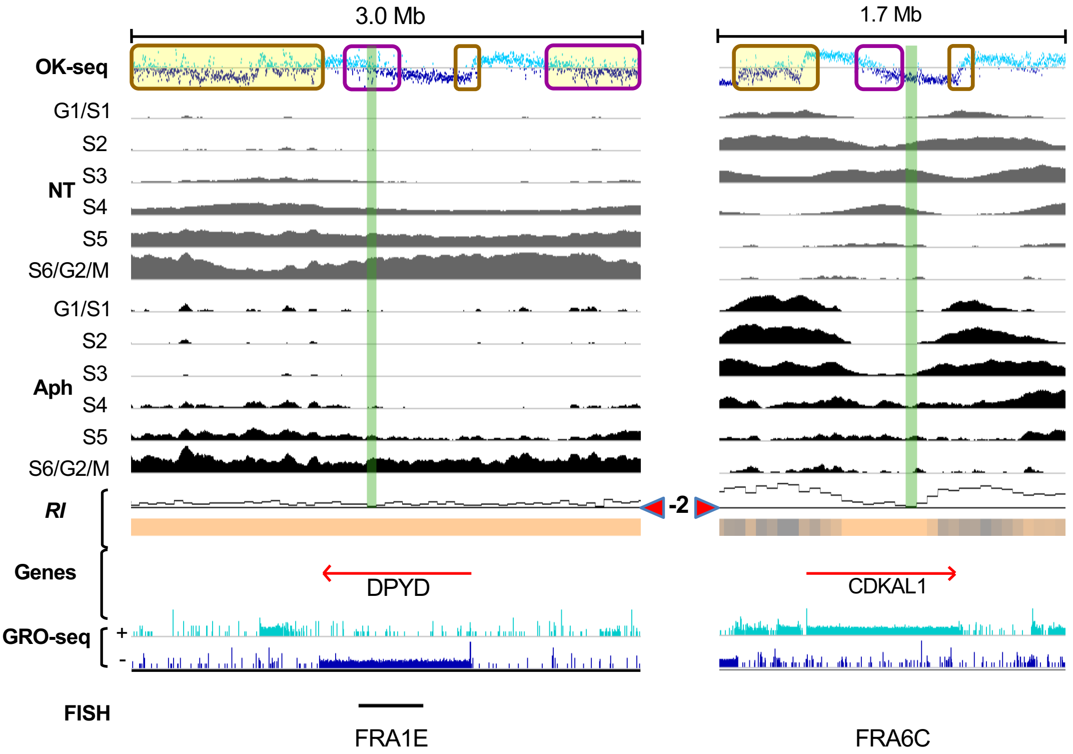

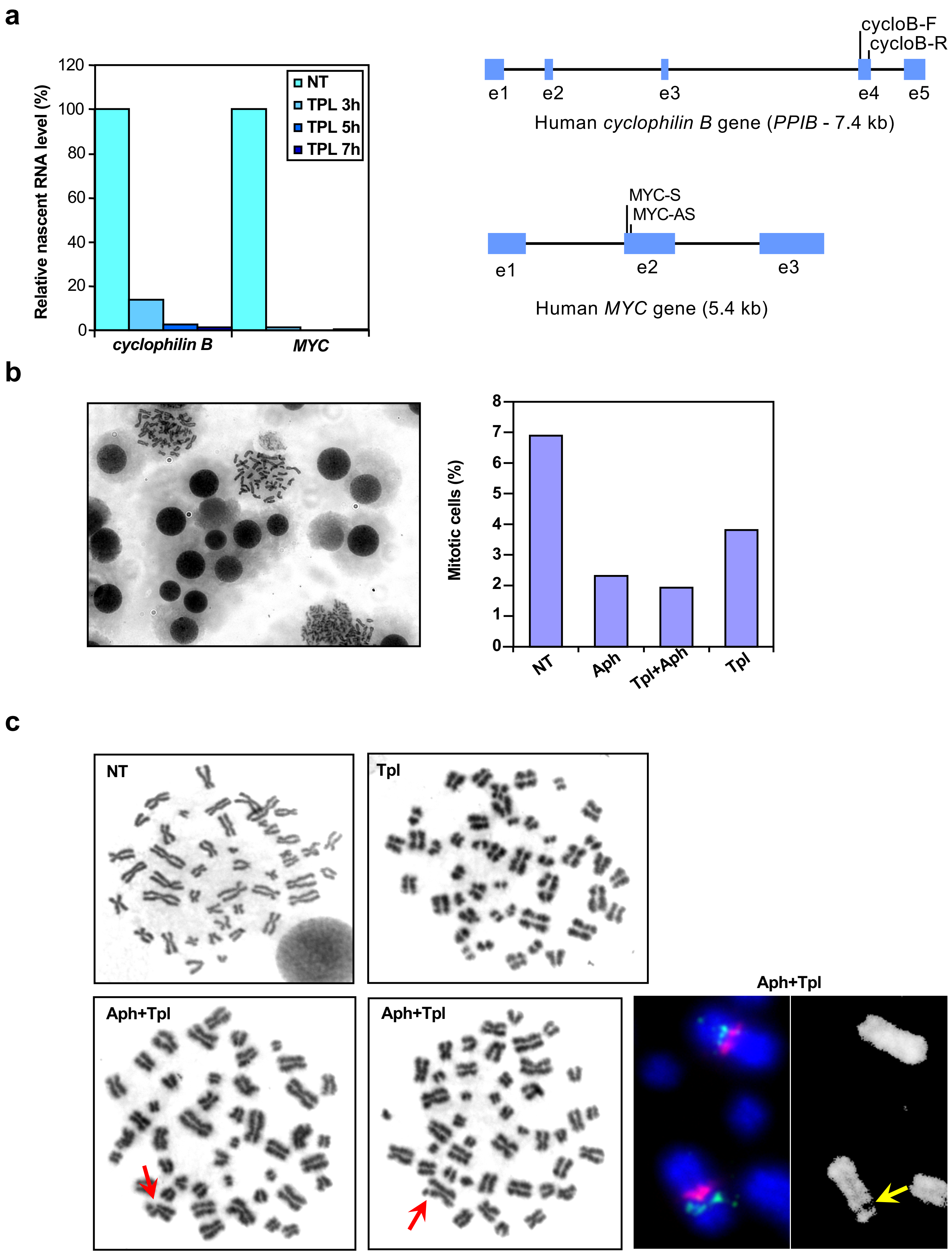
